# Uric acid lowering treatment alleviates perivascular carotid collar placement induced neointimal lesions in *Uricase* knockout mice

**DOI:** 10.1101/366096

**Authors:** Jie Lu, Ming-Shu Sun, Xin-Jiang Wu, Xuan Yuan, Zhen Liu, Xiao-Jie Qu, Xiao-Peng Ji, Tony R Merriman, Chang-Gui Li

## Abstract

Hyperuricemia (HU) is a cause of gout. Clinical studies show a link between HU and cardiovascular disease. However, the role of soluble serum urate on atherosclerosis development remains elusive. We aimed to use a new HU mouse model (*Uricase/Uox* knockout (KO)) to further investigate the relationship between HU and atherosclerosis. Mouse model of induced carotid atherosclerosis was established in the novel spontaneous HU *Uox-*KO mouse and their wild type littermates (C57BL/6J background). Mice were implanted with a perivascular collar placement around the right carotid artery in combination with a western-type diet. To investigate urate-lowering treatment (ULT) effects on intima, the mice were gavaged daily from the age of 6 weeks with allopurinol. Human umbilical vein endothelial cells (HUVECs) were co-incubated with soluble urate, with and without probenecid, to study the mechanism of urate-related atherosclerosis. The *Uox-*KO mice had significantly elevated serum urate levels combined with higher blood urea nitrogen and serum creatinine. Western blot analysis showed enhanced levels of atherosclerosis inflammatory response proteins. However, there were no other risk indicators for the pathogenesis of atherosclerosis, including increased fasting glucose, altered lipid and atherosclerosis characterized cardiovascular and histological manifestations. In contrast, collar placement *Uox-*KO mice showed severe neointimal changes in histology staining consistent with increases in intimal area and increases in proliferating cell nuclear antigen (PCNA) - and F4/80-positive cells. Allopurinol reduced neointimal areas induced by the perivascular collar in hyperuricemic mice accompanied by decreased expression of PCNA- and F4/80-positive cells (*P*< 0.05). ULT alleviated atherosclerosis inflammatory response factors and reactive oxygen species intensities in both collar placement *Uox-*KO mice and urate-stimulated HUVECs*. In vitro* results using HUVECs showed ROS was induced by urate and ROS induction was abrogated using antioxidants. These data demonstrate that urate *per se* does not trigger atherosclerosis intima lesions in mice. Urate worsens carotid neointimal lesions induced by the perivascular collar and urate-lowering therapy partially abrogates the effects. The current study warrants the further human based study on the possible benefits of urate-lowering therapy in atherosclerosis patients with HU.

**Summary statement:** We generated a carotid collar placement atherosclerosis model in the novel spontaneous HU *Uox-KO* mouse and demonstrate that urate plays a contributing rather than a causal role in the carotid neointimal lesions, while urate-lowering treatment may bring additional benefits in this HU mouse model.

## Introduction

Cardiovascular disease (CVD) is the leading cause of death world-wide (Lozano et al., 2012). Atherosclerosis is one of the major CVDs characterized by focal intimal thickening and ultimately luminal obstruction induced by fibro-proliferation and an inflammatory process mediated by cytokine production and vascular regulatory mechanisms (Zmuda et al., 2011). Neointimal formation happens at the early stage of atherosclerosis which is a complex process initiated by the damage of endothelial cells and exposure of vascular smooth muscle cells to circulating blood elements (Tian et al., 2017). Hyperuricemia (HU) causes gout and has been implicated in hypertension and atherosclerosis in humans (Odden et al., 2014). Epidemiological studies have associated HU with atherosclerotic vascular diseases (Wu et al., 2016), and the level of serum urate (SU) can predict cardiovascular outcomes and mortalities in both sexes (Xia et al., 2014). HU is also a risk factor for hypertension, which is a potent contributor of atherosclerosis (Wu and Chan, 2012). A meta-analysis of 18 prospective cohort studies, including data from more than 55,000 patients, showed an increased risk of incident hypertension in subjects with HU, and the overall risk increased by 13% per 1 mg/dL increase in SU (Grayson et al., 2011). More recently, a retrospective cohort study demonstrated that HU plays a significant role in the progression from prehypertension to hypertension with a 35% increasing ratio of hypertension risk in men (Do et al., 2015). These findings have prompted a growing research interest on the possible benefits of urate-lowering treatment (ULT) in cardiovascular diseases. However, it has not been definitively established whether urate is merely a marker or a causal agent of CVD, or whether ULT affects outcomes. This study was designed to investigate the relationship between urate and atherosclerosis in a spontaneous HU mouse model with collar-induced neointimal lesions.

## Results

### Soluble urate does not induce atherosclerotic phenotypes

SU levels in *Uox*-KO mice were almost 3-fold increased compared to WT counterparts (563.9 μmol/L ± 16.9 *vs.* 176.3 μmol/L ± 6.7, *P*< 0.001; Table 1). Compared with WT controls, BUN and serum creatinine were significantly elevated in *Uox*-KO mice (Table 1). No apparent changes occurred in the fasting glucose level of *Uox*-KO mice compared with WT mice (Table 1). Neither did lipid profiles change (TC, HDL-C and LDL-C; Table 1).

**Table 1.** Blood biochemistry in WT and *Uox*-KO mice.

|  | WT | KO | *P* value | WT-collared | KO-collared | *P* value | KO-collared +<br>allopurinol | *P* value (compared<br>to KO-collared) |
| --- | --- | --- | --- | --- | --- | --- | --- | --- |
| **Uric acid (μmol/L)** | 176.3±6.7 | 563.9±16.9 | < 0.001 | 187.5±4.9 | 573.0±16.4 | < 0.001 | 325.8±9.2 | < 0.001 |
| **Blood urea nitrogen (mmol/L)** | 4.47±0.16 | 10.04±0.35 | < 0.001 | 5.19±0.28 | 10.42±0.42 | < 0.001 | 7.91±0.63 | 0.005 |
| **Creatinine (μmol/L)** | 16.53±1.67 | 23.63±2.18 | 0.04 | 15.36±1.83 | 30.5±0.97 | < 0.001 | 21.77±2.47 | 0.0047 |
| **Fasting glucose (mmol/L)** | 6.0±0.2 | 6.3±0.2 | 0.25 | 6.1±0.2 | 6.2±0.2 | 0.71 | 6.4±0.3 | 0.6 |
| **Total cholesterol (mmol/L)** | 3.36±0.12 | 3.67±0.13 | 0.1 | 3.59±0.08 | 3.85±0.12 | 0.08 | 3.62±0.14 | 0.23 |
| **Triglycerides (mmol/L)** | 1.49±0.16 | 1.13±0.03 | 0.03 | 1.35±0.07 | 1.23±0.03 | 0.13 | 1.27±0.06 | 0.62 |
| **HDL cholesterol (mmol/L)** | 1.61±0.13 | 1.74±0.05 | 0.35 | 1.60±0.05 | 1.80±0.08 | 0.06 | 1.59±0.13 | 0.21 |
| **LDL cholesterol (mmol/L)** | 0.49±0.07 | 0.55±0.04 | 0.44 | 0.48±0.06 | 0.55±0.06 | 0.43 | 0.50±0.03 | 0.44 |

Biochemical indicators in WT and *Uox*-KO mice (*n* = 8 per group, males, 10-16 weeks of age). Data were shown as mean ± SEM.

To evaluate the effects of urate on the cardiovascular system, blood pressure, endothelium-dependent vasodilatation and cardiac function were tested. Systolic blood pressure (SBP), diastolic blood pressure (DBP) and mean blood pressure (MBP) exhibited no difference between *Uox*-KO mice and WT controls (Fig. 1A). Neither did aortae dilation nor cardiovascular function change (Fig. 1B; Table 2). This does not support a direct role of urate in the control of blood pressure, vasodilation and cardiac function. As shown in Fig 1C and 1D, pathogenic proteins in atherosclerosis were up-regulated in both plasma and carotid lesions - MCP-1, ICAM-1, and VCAM-1 (*P*< 0.05). However, *Uox*-KO mice did not present an atherosclerotic phenotype, showing no changes in intimal areas, proliferation (proxied by PCNA-positive cells) and inflammation (proxied by F4/80-positive cells) in Fig. 1E and 1F.

**Table 2.**
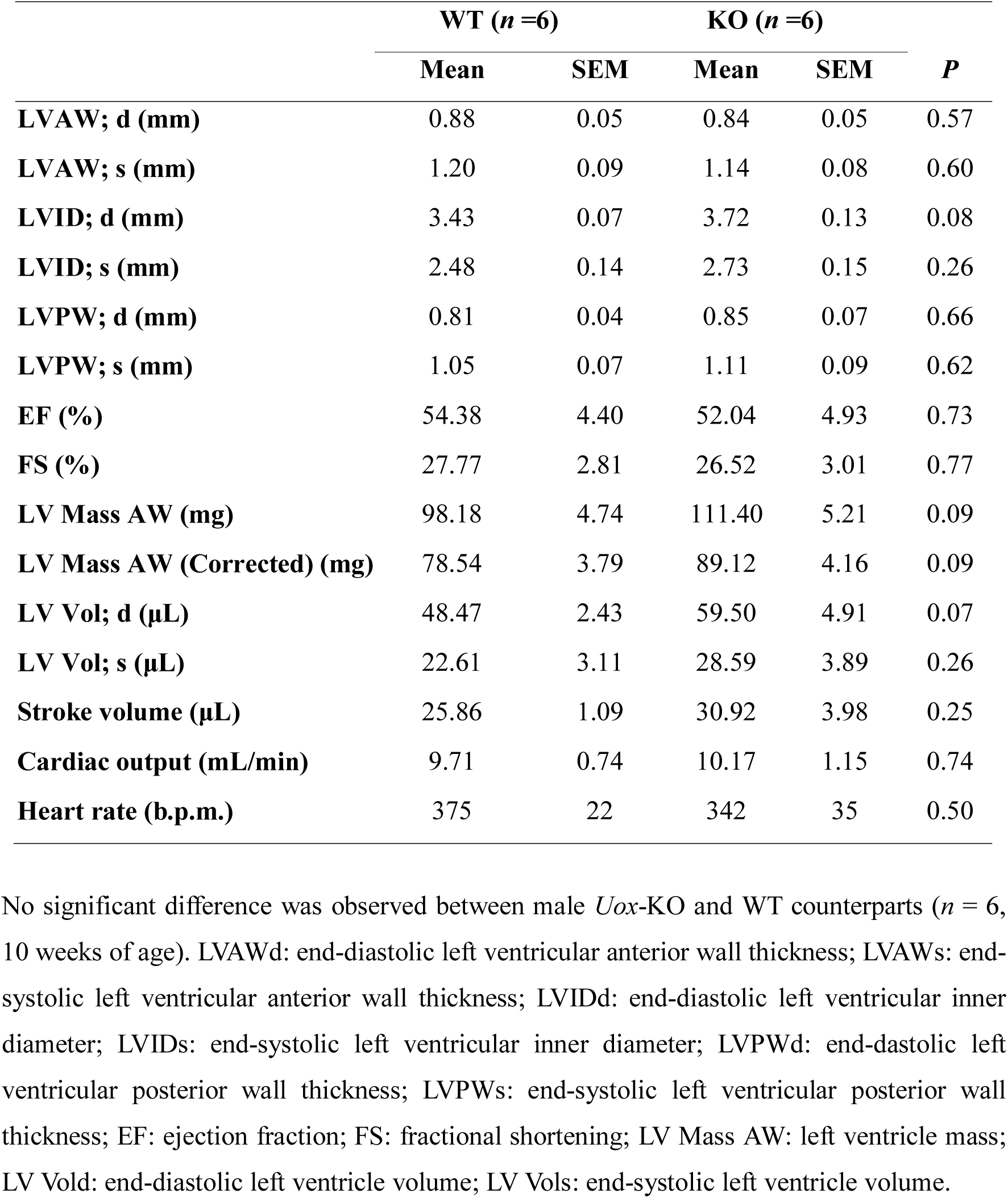
The results of transthoracic ultrasound.

**Fig. 1.**
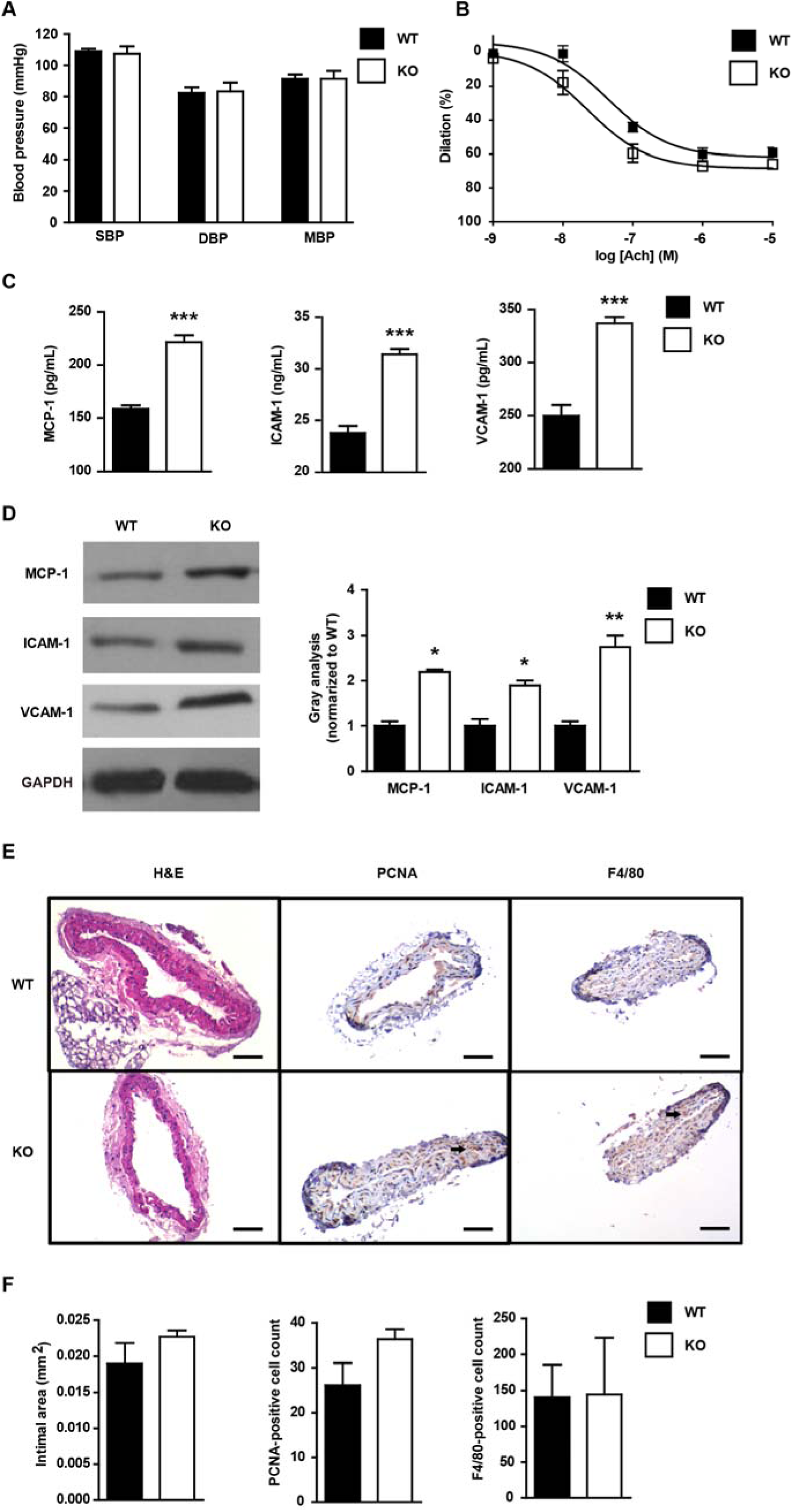
Soluble urate does not induce atherosclerosis phenotypes. **A**, Systolic (SBP), diastolic (DBP) and mean (MBP) blood pressure in non-fasting 10-week-old WT and *Uox*-KO mice (*n* = 10, 7). **B**, Endothelium-dependent vasodilatation of aortic segments in 10-week-old WT and *Uox*-KO mice (*n* = 3) induced by acetylcholine (Ach). **C,** Protein levels of MCP-1, ICAM-1, and VCAM-1 in serum of 10-week-old mice (*n* = 6) detected by ELISA. **D,** Protein levels of MCP-1, ICAM-1, and VCAM-1 in carotid arteries of 10-week-old mice determined by three independent western blotting experiments (*n* = 3). **E and F**, Carotid pathological assessment in WT and *Uox*-KO mice (*n* = 6) quantified by intimal area, PCNA-and F4/80-positive cell counts. Bars = 50μm.^*^*P*＜0.05, ^**^*P*＜0.01, ^***^*P*＜0.001 versus WT. Error bars represent SEM.

### Hyperuricemia accelerates carotid neointimal lesions with collar placement

We next addressed the questions: is the effect of urate evident only once a stress is given, and does ULT affect the atherosclerosis pathogenesis? This perivascular carotid collar model along with western-type diet represents a useful complementary model in which neointimal lesions show up at the early-stage of atherosclerosis. With collar induction HU makes neointimal lesions worse, indicated by the increases in intimal area, more PCNA- and F4/80-positive cells in the carotid artery compared with WT collared mice (Fig. 2C, 2D). Administration of allopurinol (100 mg/kg), a xanthine oxidase inhibitor and urate-lowering drug, significantly alleviated the intimal phenotypes including intimal area, PCNA-positive cell counts and F4/80 positive cell counts. Allopurinol also alleviated both SU and renal functions indicated by BUN and creatinine in *Uox*-KO collared mice compared to WT collared mice (Table 1). Consistently, allopurinol decreased expression of MCP-1, as well as ICAM-1 and VCAM-1, in both plasma and carotid tissues determined by ELISA and RT-PCR, respectively (Fig. 2A, 2B). The down-regulation of MCP-1, ICAM-1, and VCAM-1 in carotid tissue indicated that inflammation was improved by ULT. Echocardiographic analysis of heart rate, diastolic left ventricular internal diameter and ejection fraction exhibited no difference between collared *Uox*-KO mice and WT controls (data not shown).

**Fig. 2.**
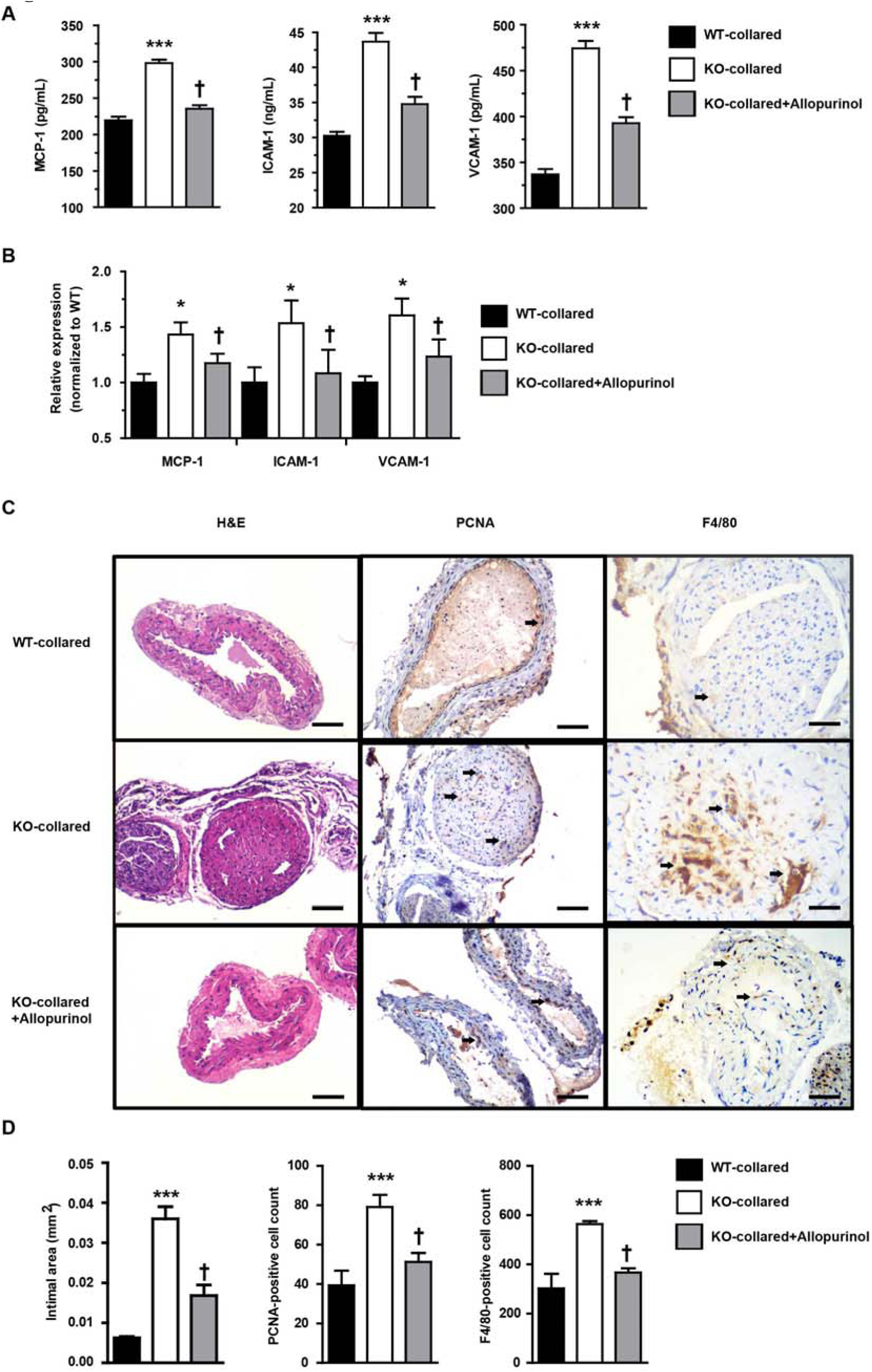
Hyperuricemia accelerates carotid neointimal lesions with collar placement. Shown are effects of collar induction and 10-week allopurinol treatment (100 mg/kg) on MCP-1, ICAM-1, and VCAM-1 levels in **A** plasma (*n* = 6) and **B** carotid tissues (*n* = 6), and carotid morphology, proliferation and inflammation represented by intimal area, PCNA-and F4/80-positive cell counts in **C and D** (*n* = 6), respectively. Bars = 50μm. ^***^*P*＜0.001 versus WT control and ^†^*P*＜0.05 versus untreated *Uox*-KO mice. Error bars represent SEM.

### Hyperuricemia elevates ROS in carotid artery *in vivo*

Given ROS plays a putative role in the pathogenesis of atherosclerosis, we tested the ROS intensity by fluorescent dye dihydroethidium staining in the carotid artery of WT and *Uox*-KO mice. *Uox*-KO mice had elevated ROS levels compared with WT, with or without collar placement (Fig 3A, 3B). ROS intensities were reduced in *Uox*-KO mice with 100 mg/kg allopurinol treatment for 10 weeks *versus* WT controls in the presence or absence of collar placement (Fig 3A, 3B), implying that urate imposes additional oxidative stress which may accelerate atherosclerosis development.

**Fig. 3.**
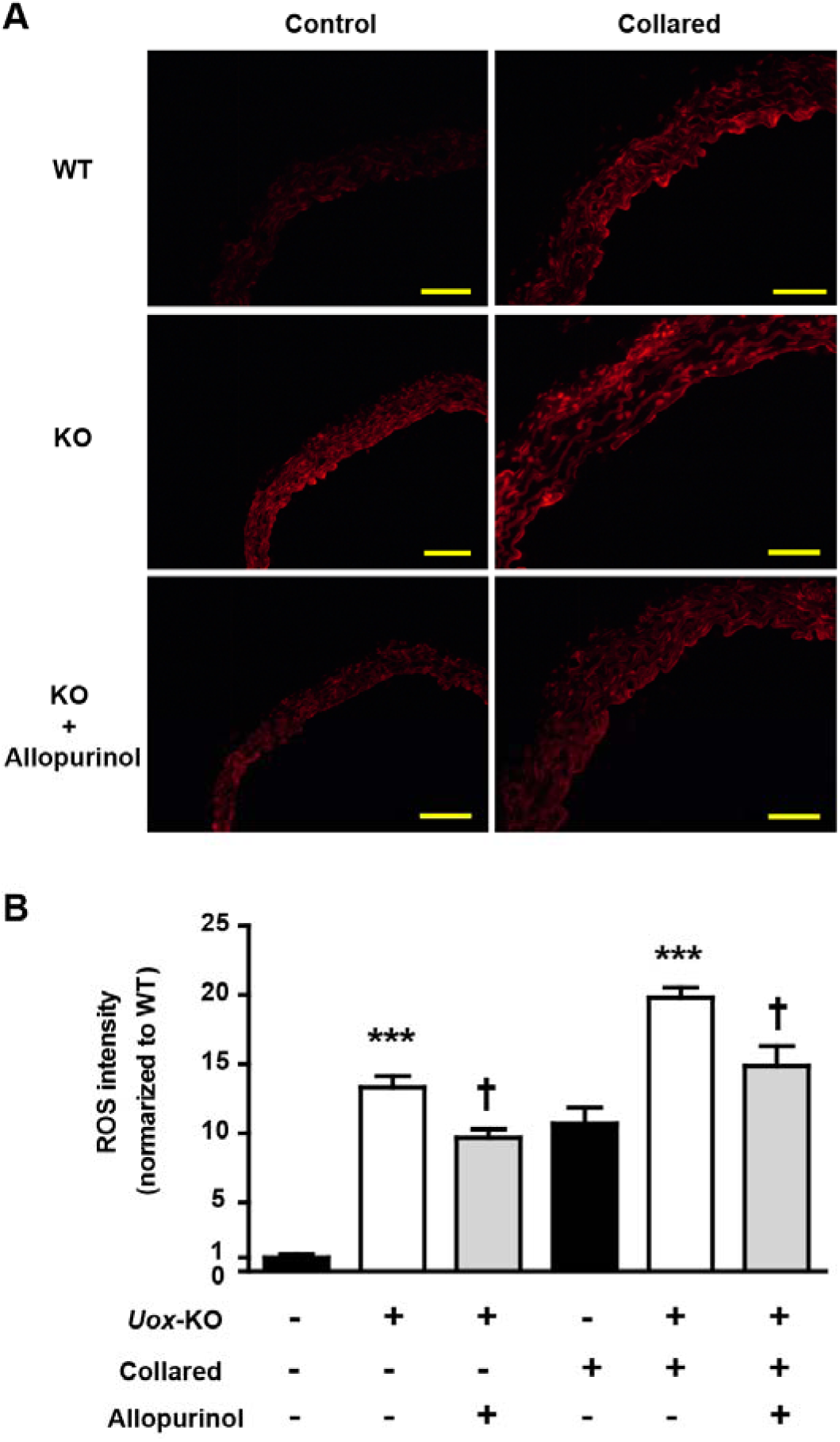
Hyperuricemia elevates ROS in carotid artery in vivo. **A**, Representative images of fluorescent dye dihydroethidium staining from carotid artery of WT and *Uox*-KO mice (*n* = 6). **B,** Effects of 8-week allopurinol treatment (100 mg/kg) on ROS were quantified. Bars = 50μm. ^***^*P*＜0.001 versus WT control. ^†^*P*＜0.05 versus untreated *Uox*-KO mice. Error bars represent SEM.

### Soluble urate induces ROS enhancement *in vitro*

Human umbilical vein endothelial cell (HUVEC) (10^5^/well) viability was measured when co-incubated with soluble urate (200 μmol/L, 400 μmol/L, 600 μmol/L and 800 μmol/L) at 24h, 48h and 72h. As shown in Fig. 4A to 4C, soluble urate decreased cell viability in a dose-dependent and time-dependent manner and elevated ROS levels – for example with 800 μmol/L urate, ROS intensity was almost 10-fold increased. ROS levels were extenuated by pre-incubation for 12h with probenecid (1 mmol/L), an organic anion transport inhibitor, or N-acetylcysteine (NAC) (10 μmol/L), a ROS scavenger, or with both (Fig. 4D). Relative mRNA expression of MCP-1, ICAM-1, and VCAM-1 in soluble urate (800 μmol/L, 48h) treated HUVECs was significantly higher than controls and this was lessened by pre-incubation with probenecid or NAC or both (Fig. 4E).

**Fig. 4.**
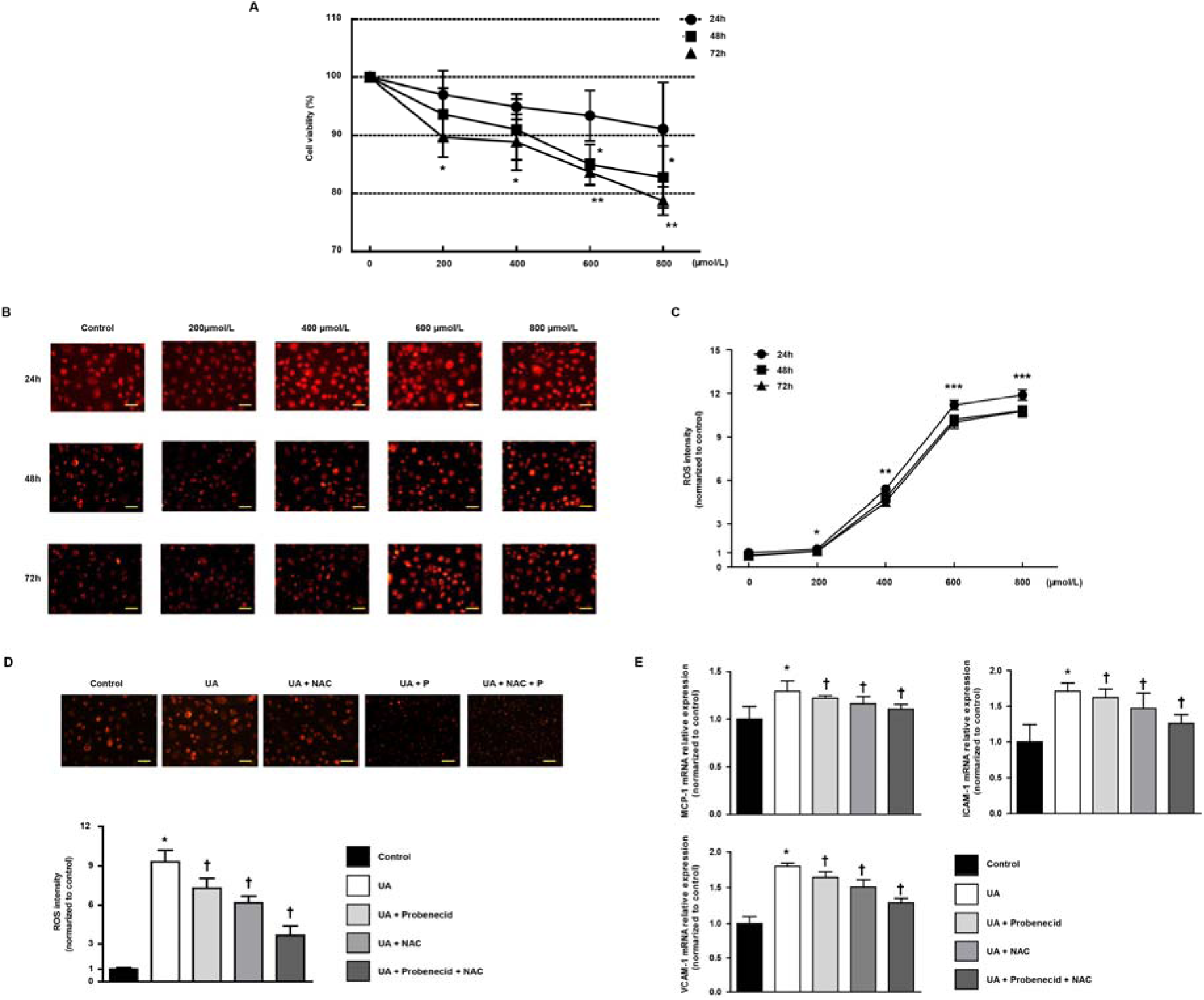
Soluble urate induces ROS enhancement in vitro. **A,** HUVECs (10^5^/well) viability measurement co-incubated with soluble urate (200, 400, and 800 μmol/L) at 24h, 48h and 72h. **B and C,** Representative images of fluorescent dye dihydroethidium (DHE) staining from soluble urate stimulated HUVECs and quantified ROS levels. **D,** DHE staining on soluble urate (800 μmol/L, 48h) stimulated HUVECs after 12h pre-incubation with 1 mmol/L probenecid or 10 μmol/L N-acetyl-l-cysteine (NAC) or both. **E,** Relative mRNA expression of MCP-1, ICAM-1, and VCAM-1 in HUVECs. Data were represented as mean ± SEM. Bars = 20μm.^*^*P*＜0.05, ^**^*P*＜0.01, ^***^*P*＜0.001 versus control, ^†^*P*＜0.05 versus UA group. Error bars represent SEM.

## Discussion

The presence of hepatic *Uox* is the reason that rodents have lower SU compared to humans (Alvarez-Lario and Macarron-Vicente, 2010). When *Uox* is knocked out, mice develop spontaneous HU similar to the SU level of humans (Lu et al., 2018). Based on this HU mouse model, we further generated an atherosclerosis model with right carotid artery peri-collar placement and a western-type diet. Using this model we showed that, because *Uox*-KO mice did not present an atherosclerotic phenotype despite of facts that inflammatory response and ROS indeed were induced in *Uox*-KO mice. However, HU worsened the development of neointimal lesions through *in vivo* ROS enhancement.

Epidemiological studies have detected an association between HU and hypertension, while evidence for causation, that includes data from Mendelian randomization studies, is limited and inconclusive (Dehghan et al., 2008; Sluijs et al., 2015; Vitart et al., 2008). Hypertension is a risk factor in the pathogenesis of atherosclerosis. Here we describe the cardiovascular characteristics in *Uox*-KO male mice, which exhibit no signs of heart dysfunction, heart morphology alteration or blood pressure change. This is consistent with data from another HU mouse model with liver *Glut9* deficiency (Preitner et al., 2015). These data therefore do not support a direct causal role of urate on blood pressure in mice.

To our knowledge, this study is the first to assess conducted dilation responses of aortae in the spontaneous HU mouse and showed that urate cannot induce experimentally significant changes of endothelium-dependent vasodilatation. Indeed, when expressed as a function of the dilation, the conducted dilation of aortae in *Uox*-KO and WT mice was relatively similar. Interestingly, the endothelial relaxation dysfunction reported in aortae from *low-density lipoprotein receptor/apolipoprotein E* (*LDL/ApoE*) double knockout mice was in regions with significant lesions, and not in other regions or in aortae from the *ApoE* single KO mice where lesions were minimal (Beleznai et al., 2011). Similarly, even in diabetic *ApoE*-KO mice, endothelial dysfunction was only reported in plaque-prone regions of the aortae, while plaque-resistant segments maintained a normal acetylcholine response (Ding et al., 2005). These pieces of evidence are consistent with our observation of no vasodilation dysfunction in our *Uox*-KO mice compared with WT mice.

Our results show that spontaneous HU without any stress does not induce obvious atherosclerosis phenotypes. However, hyperuricemia alone was sufficient to induce oxidative stress and enhance levels of atherosclerosis associated inflammatory cytokines indicating that HU is only a promoting factor rather than an initiator in atherosclerosis. HU may contribute to an activated atherosclerotic pathological process. MCP-1, ICAM-1 and VCAM-1, crucial pathogenic elements in atherosclerosis, are up-regulated in atherosclerotic lesions and influence growth factor production and medial smooth muscle cell migration (Dzau et al., 2002; Hsueh et al., 2016). HU enhanced MCP-1, ICAM-1 and VCAM-1 as shown in serum level and protein expression, suggesting that these three molecules are also involved in a HU-driven atherosclerosis promoting effect that would exacerbate the pathogenesis of atherosclerosis. The augmented response of MCP-1, ICAM-1 and VCAM-1 to a given concentration of urate would stimulate downstream inflammation and thereby induce atherosclerosis progressing to a greater extent than controls *in vitro*. Increasing of intimal area due to smooth muscle cell movement and reproduction is an essential component of atherosclerosis, which can be indicated by proliferating cell nuclear antigen (PCNA) (Lu et al., 2017). Macrophages accumulate in atherosclerotic plaques, playing crucial roles in atherosclerotic immune responses (Li et al., 2017). Significant increases in intimal areas, PCNA- and F4/80-positive cells in the carotid artery indicates that HU contributes to atherosclerosis, though no plague were found. Despite a few hundred systematic reviews, meta-analyses, and Mendelian randomisation studies exploring 136 unique health outcomes, convincing evidence of a clear causal role of urate in disease pathogenesis only exists for gout and nephrolithiasis (Liu et al., 2015). Urate is involved in a diverse array of biological functions, while possibly contributing to the pathogenesis of cardiovascular phenotypes, rendering it a pathogenic but not causal role (Alvarez-Lario and Macarron-Vicente, 2010). HU did associate with an increased risk of cardiovascular death only in participants with gout and existing cardiovascular disease (Muoio and Newgard, 2008), consistent with our experimental *in vivo* data.

Reactive oxygen species are another major but non-specific mediator in the formation of atherosclerosis-inducing endothelial dysfunction. It can reduce the bioavailability of nitric oxide, a potential anti-atherosclerotic factor (Montezano and Touyz, 2012). Urate can lead to ROS enhancement which facilitates atherosclerosis by oxidative stress (Forstermann et al., 2017), which is consistent with our *in vivo* and *in vitro* data. Changes in ROS exhibited a similar trend as the change of neointimal lesions in collar-induced *Uox*-KO mice, which were partially rescued by ULT. This suggests that ROS may also contribute to urate-induced atherosclerosis-promoting effects. The effect of allopurinol (via the active metabolite oxypurinol) that inhibits xanthine oxidase (XO) activity and suppresses urate biosynthesis (Elion, 1966), also reduces ROS production by inhibiting XO. Other non-XO effects of allopurinol include LDL oxidation prevention, heat shock protein expression inhibition and decreasing early changes in inflammation such as leukocyte activation by reducing adherence, rolling and extravasation (George and Struthers, 2009). Given our data shows HU was sufficient to induce both ROS production and atherosclerosis associated inflammatory cytokines, worsened ROS level, inflammatory molecules and neointimal lesions observed in collared *Uox*-KO mice could be effects due to an “additional action” imposed by HU. Thus, all the neointimal lesions and associated inflammatory factors as well as ROS production were alleviated when ULT was present. *In vitro*, our study showed the same trend of ROS *in vivo*. ROS intensities were lowered with probenecid intervention. Combined with probenecid and NAC, the strongest ROS lowering effect was observed (in HUVECs). Thus, the benefits of XO inhibitors such as allopurinol might rely on blocking the production of oxidants rather than on lowering urate (Johnson et al., 2013). A random, double-blind, crossover study also showed that the mechanism of improvement in endothelial function with high-dose allopurinol lies in its ability to reduce vascular oxidative stress and not in urate reduction (Civelek and Lusis, 2014). Therefore, although our data exclude a direct causal role of urate *per se* on atherosclerosis in *Uox*-KO mice, further studies to address a potential role of XO activity on the cardiovascular function are warranted. Overall, this work reinforces the conclusion that urate accelerates pathogenesis of atherosclerosis and ROS lowering may bring the anti-atherosclerotic effects.

It remains controversial as to whether asymptomatic HU should be treated for the purpose of improving cardiovascular outcomes (Abeles, 2015). Stamp and Dalbeth (Stamp and Dalbeth, 2017) suggested that asymptomatic HU treatment needs to be cautiously considered, due to the limited data and the potential risks of treatment. Kok *et al.* (Kok et al., 2014) reported that allopurinol therapy in patients with gout does not yield beneficial cardiovascular outcomes. However, Kuwabara *et al*. (Kuwabara et al., 2017) promoted the use of ULT for asymptomatic HU in a five-year Japanese cohort study with 13,201 subjects. ULT was associated to better outcomes in HU patients with cardiovascular diseases accompanied by benefits in endothelial dysfunction and systemic inflammation (Hayashino et al., 2016). A randomised controlled trial reported that allopurinol reduces central blood pressure and carotid intima-media thickness progression at 1 year in patients with recent ischemic stroke and transient ischemic attack (Higgins et al., 2014). A mouse study reported that allopurinol represents a potential novel strategy for preventing left ventricular remodelling and dysfunction after myocardial infarction (Engberding et al., 2004). As well, significant anti-atherosclerotic effects were seen in a HU study in *ApoE*-KO mice (Wakuda et al., 2014). Consistently, our results suggest that ULT with allopurinol for hyperuricemic mice may improve neointimal lesions significantly.

*Uox-*KO mice did not show significant difference in blood pressure, vasodilation or atherosclerotic lesions as compared with controls even urate concentration >560 μmol/L, comparable to human (Feig et al., 2008). Even though the *Uox-*KO mice developed chronic renal dysfunction with a trend for increased plasma BUN, creatinine and chronic renal inflammation and fibrosis (Lu et al., 2018), there was no correlation between renal or metabolic dysfunction and atherosclerosis development in this mouse model.

It is important to note that there was only approximately 40% of survivors in the *Uox-*KO birth cohort (Lu et al., 2018). Thus, the survived *Uox*-KO mice with constant high UA levels may represent biased subjects and they normally develop significant metabolic and renal dysfunction, which actually mimics the human cases of gout and hyperuricemia with severe complications. A transient increase in uricosuria during the first two weeks of life in *Uox*-deficient mice might be responsible for the relative low survival rate (Flannick and Florez, 2016). The current model is useful to study the asymptomatic HU from a metabolic perspective though it is not fully understood the reasons of poor survival rate.

There is phenotypic heterogeneity existed between sexes in the *Uox*-KO model. Female *Uox*-KO mice have elevation in blood pressure compared with WT counterparts, while males do not (Lu et al., 2018). In addition, the association between SU and hypertension might vary according to age and sex, being more significant in younger and female subjects (Batzoglou and Schwartz, 2014). To avoid confounding factor due to gender or sex hormone difference, we only chose male mice in the current study. Further investigation in females would help to delineate a full picture of the effect of HU on cardiovascular diseases.

In conclusion, this is the first evidence to demonstrate that urate plays a contributing rather than a causal role in the carotid neointimal lesions, while ULT may bring additional benefits in this spontaneous HU mouse model. Clinical and mechanism studies are warranted to investigate the ULT’s anti-atherosclerotic benefits in atherosclerosis patients with HU.

## Materials and Methods

### Mouse model

The spontaneous HU mice were developed by knock-out of the hepatically-expressed *Uricase* (*Uox*) gene (Lu et al., 2018). Controls were their wild type (WT) littermates on the C57BL/6J background. As previously described (Baetta et al., 2007), we generated perivascular carotid collar placement mouse models with non-occlusive silastic collars (length, 5 mm; internal diameter, 0.3 mm; external diameter, 0.6 mm) in 10-week-old males. Four weeks before surgery, the animals received high-fat, high-cholesterol (HF/HC) western-type diets (21% (wt/wt) fat and 0.15% cholesterol). Mice were anesthetized by ketamine/xylazine and kept on continuous anesthesia during the surgery. Subsequently, a midline neck incision was made to surgically expose the right carotid artery, and the collar was positioned around the right carotid artery and held in place with a nylon sleeve. Both carotid sheaths were openned, and the common carotid arteries were dissected free from the surrounding connective tissue. The carotid arteries were then returned to their original position and the wound was sutured. After a 6-week collar placement with western-type diet, all mice were sacrificed for subsequent measurements. To evaluate the urate-lowering effect, mice were gavaged with allopurinol at a dose of 100 mg/kg from 6 weeks to 16 weeks of age with collar induction at 10-week-old.

Mice were housed under specific pathogen free (SPF) conditions at 22°C under a 12-h light/dark photoperiod with *ad libitum* access to rodent diet and sterile water (Zhang et al., 2010). The Animal Research Ethics Committee of the Affiliated Hospital of Qingdao University approved this study.

### Blood biochemistry

Mice were fasted overnight before blood collection from the outer canthus. All biochemical indicators including SU, blood urea nitrogen (BUN), serum creatinine, fasting glucose, lipid profiles (total cholesterol (TC), triglyceride (TG), high- and low-density lipoprotein cholesterol (HDL-C, LDL-C) were measured by an automatic biochemical analyser (Toshiba).

### ELISA

Plasma monocyte chemoattractant protein-1 (MCP-1), intercellular adhesion molecule-1 (ICAM-1) and vascular cell adhesion molecule-1 (VCAM-1) levels were quantified using an enzyme-linked immunosorbent assay (ELISA) kit (R&D Systems).

### Quantitative real-time RT-PCR

Total RNA was isolated from the carotid artery or cell line using trizol reagent (Roche Pharmaceuticals) and then reverse transcribed using a Fast Quant RT kit (Takara-Bio), followed by amplification using primers for MCP-1, ICAM-1 and VCAM-1 under the condition: 95°C for 10 min and 40 cycles of 95°C for 15s, 58°C for 20 s and 68°C for 20 s. The threshold cycle (Ct) was determined and used to calculate ΔCT values. The ΔΔCt (2^−ΔCt^) was used to calculate relative mRNA expression, with each measurement performed in triplicate. Primer sequences are shown in Supplementary Table 1.

### Western blot

Protein was extracted from the carotid artery. 50 μg protein lysates were loaded and transferred to nitrocellulose membranes. Blots were incubated with the primary antibodies (1:2000, Abcam) against MCP-1 (Catalog no. ab25124), ICAM-1 (Catalog no. ab119871), VCAM-1 (Catalog no. ab134047) and GAPDH (Catalog no. ab8245) at 4°C overnight. After incubation with horseradish peroxidase-conjugated goat anti-rabbit secondary antibodies, visualization was performed with an enhanced chemiluminescence kit (Thermo Scientific). Each measurement was performed in triplicate.

### Echocardiographic analysis

Echocardiography was performed after anesthesia with isoflurane using a Vevo 2100 ultrasound system equipped with a MS400 probe (VisualSonics) for *in vivo* transthoracic ultrasound imaging. The heart was imaged in a 2-dimensional mode in the parasternal long-axis view. An M-mode cursor was positioned perpendicular to the interventricular septum and the posterior wall of the left ventricle at the level of the papillary muscles. Stroke volume was calculated as (LV Vol; d – LV Vol; s) and cardiac output as ([LV Vol; d – LV Vol; s] × HR) / 1000.

### Blood pressure

Systolic (SBP) and diastolic (DBP) blood pressure was measured by the CODA programmable noninvasive tail-cuff sphygmomanometer (Kent Scientific). Mice underwent an acclimation period of 7 consecutive days to the sphygmomanometer before experiments. Mean blood pressure (MBP) was calculated as (DBP + 1/3 [SBP – DBP]).

### Assessment of endothelium-dependent vasodilatation

Vasorelaxation of isolated aortic ring segments were determined in oxygenated Kreb’s solution. Aortic rings were precontracted with 0.1 μmol/L noradrenaline after an equilibration period of 60 min. Dilation at each acetylcholine (0.001, 0.01, 0.1, 1.0, 10 μmol/L) concentration was measured and expressed as the percentage in response to noradrenaline.

### Pathology analysis and immunohistochemistry

Carotid arteries were removed and fixed in formalin followed by paraffin embedding of 5 μm serial sections. Tissue serial sections were incubated with anti-mouse proliferating cell nuclear antigen (PCNA, 1:50, Santa Cruz, Catalog no. sc25280) and anti-mouse F4/80 (macrophage-specific marker, 1:50, Santa Cruz, Catalog no. sc52664) rabbit polyclonal antibodies. Images were captured by Nikon Eclipse TE2000-S microscope (Nikon) and analysed by Image-Pro Plus software (version 6.0).

### Reactive oxygen species (ROS) measurement

Samples were incubated with 2 μmol/L dihydroethidium (DHE) fluorescence probe (Thermo Scientific) for 30 min at 37°C in the dark to measure ROS levels. Fluorescence was determined using a Nikon 90i (Nikon) with excitation wavelength at 480 nm and emission wavelength at 610 nm.

### Soluble urate

As previously described (Eisenbacher et al., 2014), soluble urate was prepared by dissolving uric acid (UA, Sigma) in warmed media containing 1 M NaOH. The solution was tested to be free of mycoplasma, endotoxin and filtered before use. Crystals were not detectable under polarizing microscopy, nor did they develop during cell incubation.

### Cell Culture

Human umbilical vein endothelial cells (HUVECs, Cell bank of Chinese Academy of Sciences) were cultured in human endothelial cell-specific growth medium C-22010 (PromoCell) with 10% fetal bovine serum, 100 units/ml penicillin and 100 μg/ml streptomycin (Invitrogen). The viability of HUVECs was measured by Cell Counting Kit-8 (Beyotime Institute of Biotechnology) according to the manufacturer’s instruction. Cells (10^5^/well) were plated in 96-well plates and co-incubated with soluble urate (200-800 μmol/L) after 1 mmol/L probenecid (an organic anion transport inhibitor that blocks UA entry into cells) or 10 μmol/L N-acetyl-l-cysteine (NAC, a ROS scavenger) intervention for 12 hours.

### Statistical analysis

All statistical analyses were performed using GraphPad Prism software (version 7). Data were presented as the mean ± SEM. Differences between groups were analyzed by Student’s t test or one-way analysis of variance followed by Newman-Keuls multiple comparison test as appropriate. *P*< 0.05 was considered to be statistically significant.

## Acknowledgments

All authors have written and edited the article.

---

**Table 1.**
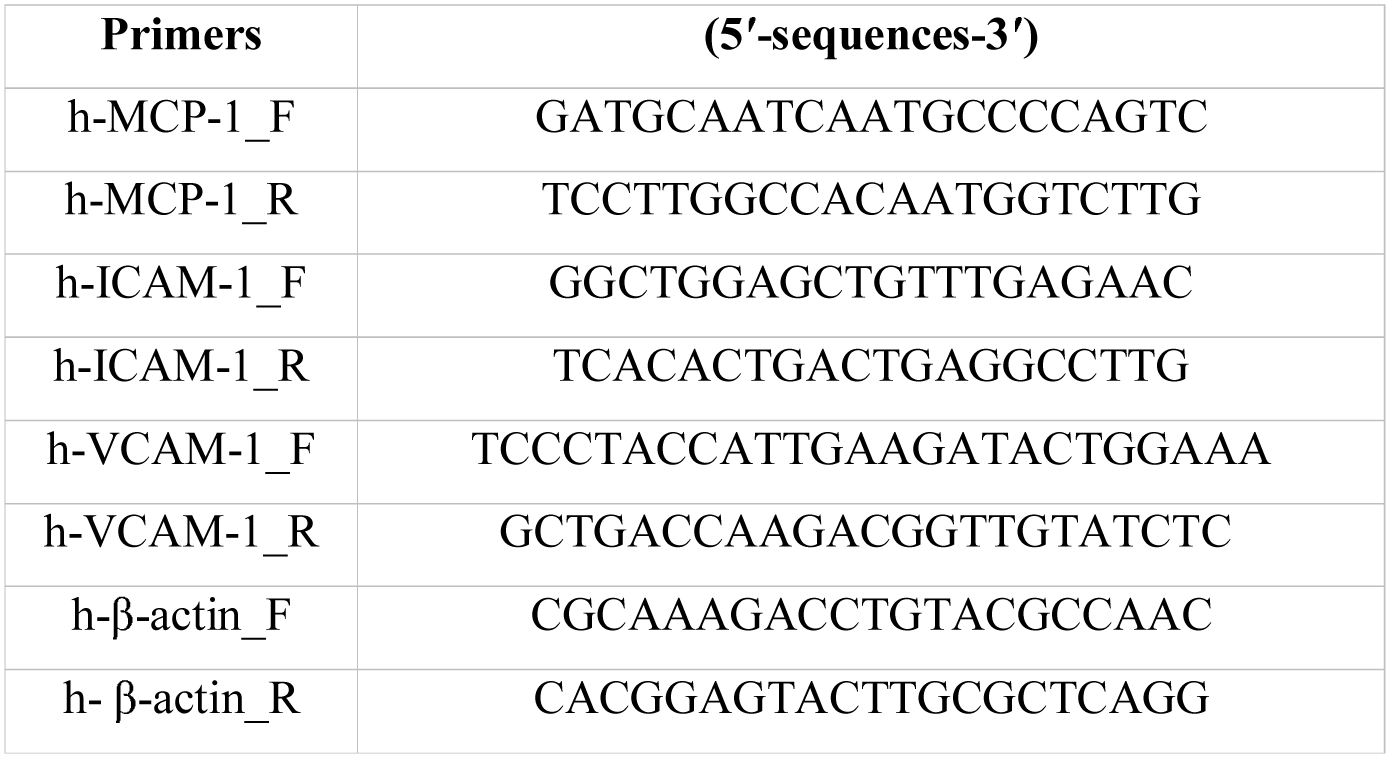
Primers of quantitative real-time RT-PCR.

